# Gestation length drives brain size and litter size variation in eutherian mammals

**DOI:** 10.1101/2024.11.13.623395

**Authors:** Thodoris Danis, Dana Lin, Daniel S. Caetano, Gregory F. Funston, Antonis Rokas

## Abstract

The length of gestation in eutherian mammals, which is key to their reproductive success, is closely connected to other life history traits and with body mass and brain mass, but causal relationships between these variables are unclear. Here, we used an integrative analytical framework to evaluate the evolutionary relationships between gestation length and eight other traits on a dataset of 3,258 eutherian mammals and infer causality. We identify variation in generation length and litter size as the primary predictors of eutherian gestation length variation, whereas additional traits, such as brain mass, significantly predict gestation length only in specific mammalian orders. Using a structural equation modeling approach known as phylogenetic path analysis to infer causality, we find that gestation length variation positively influences brain mass variation and negatively influences litter size variation. Furthermore, body mass causally influences gestation length variation only in certain orders. Consistent with these causal inferences, examination of trait-trait coevolution reinforces that gestation length is strongly positively associated with brain mass, strongly negatively associated with litter size, and only moderately correlated with body mass. These findings reveal how gestation length directly and indirectly influences, and is influenced by, other key eutherian traits. Our study establishes a robust framework for identifying causal relationships within suites of correlated and co-evolving traits.

## Introduction

The diversity in gestation duration or gestation length among eutherian mammals is substantial, varying from a few weeks in certain rodents to nearly two years in elephants [1–6]. Understanding variation in gestation length and its relationship with other traits is crucial for understanding the complex and multifaceted life history patterns of eutherian mammals. Gestation length is a critical determinant of reproductive success and offspring survival [5,7] and has previously been shown to be correlated with a variety of traits, such as body mass, brain mass, and litter size [1,2,5,6,8–13]. Although the links between gestation length and body mass, brain mass, and litter size have been well studied, the relationship between gestation length and the diversity of (potentially correlated and coevolving) mammalian life history traits remains understudied [1,2,4,9,12,14].

Prior studies that directly examined relationships between gestation length and other traits have often been limited by smaller sample sizes, with the data heavily skewed toward well-studied orders such as primates, artiodactyls, or carnivores [2,6,9,12,14,15]. This taxonomic bias may potentially overlook distinct evolutionary patterns in other orders that are highly diverse and/or have distinct ecologies, such as rodents or bats. Additionally, many of these studies have often treated traits in isolation, failing to account for the potential interactions or coevolution between multiple life history traits, which can obscure more nuanced relationships between gestation length and other traits. For instance, focusing solely on pairwise relationships between body mass, longevity, brain mass and gestation length [2,6,9,12,15–17] may neglect the ways in which these traits interact with one another, or how they jointly affect gestation length. A key conclusion of previous work is that gestation length is correlated with many other traits both across mammals as well as in specific orders [1,2,6,12,15,16]. However, whether gestation length causally influences or is influenced by other traits remains an open question.

By analyzing data of nine traits from 3,258 eutherian mammals, we investigated the evolutionary interplay between gestation length and eight other traits across eutherian mammals and in six of the most populous mammalian orders. Using state-of-the-art statistical methodologies, including phylogenetic path [18–20], machine learning [21], and trait-trait coevolution analyses [22], we identified the mammalian traits that causally influence and are influenced by gestation length.

## Results

### Several traits, including litter size, generation length and brain mass, predict gestation length

To better understand the variation of gestation length and potentially identify traits that predict its variation, we gathered data for 3,258 eutherian mammals (**Data S1**), each of which can be readily mapped to the phylogeny from Álvarez-Carretero et al. [23]. Trait values for body mass (BM), brain mass (BrM), age of female maturity (FM), interbirth interval (II), weaning age (WA), generation length (GR), gestation length (GL), litter size (LS), and longevity (L) were retrieved from the PanTheria [24], AnAge [25], EltonTraits [26] and COMBINE [27] databases (see **Methods**). Analyses were performed on seven datasets: the full dataset (all eutherian mammals; *n*=3,258 species) and datasets for six of the most speciose mammalian orders (Rodentia, *n*=1,326; Chiroptera, *n*=709; Primates, *n*=285; Eulipotyphla, *n*=258; Artiodactyla (excluding Cetacea), *n*=198; and Carnivora (excluding Pinnipedia), *n*=197). To prevent skewed results and maintain focus on the evolution of gestation length in terrestrial mammals, we excluded Cetacea and Pinnipedia due to their small sample sizes and unique aquatic adaptations, which significantly impact their life history traits and physiology [28,29].

To predict variation in gestation length based on patterns of variation in the other eight traits, we trained a Random Forest machine learning algorithm on all seven datasets (**Fig. 1**, **Table S1; Methods**). We found that generation length and litter size were the two traits that best predicted gestation length across all eutherian mammals, followed by brain mass, and body mass. Notably, the best predictors of gestation length often differed across orders; brain mass was the trait that best predicted gestation length in Rodentia, Chiroptera, and Artiodactyla, female maturity in Primates, generation length in Eulipotyphla, and body mass in Carnivora. Furthermore, while litter size and weaning age were not the best predictors in any dataset, litter size was among the top three predictors of gestation length in six datasets, and weaning age in four datasets.

**Fig 1.**
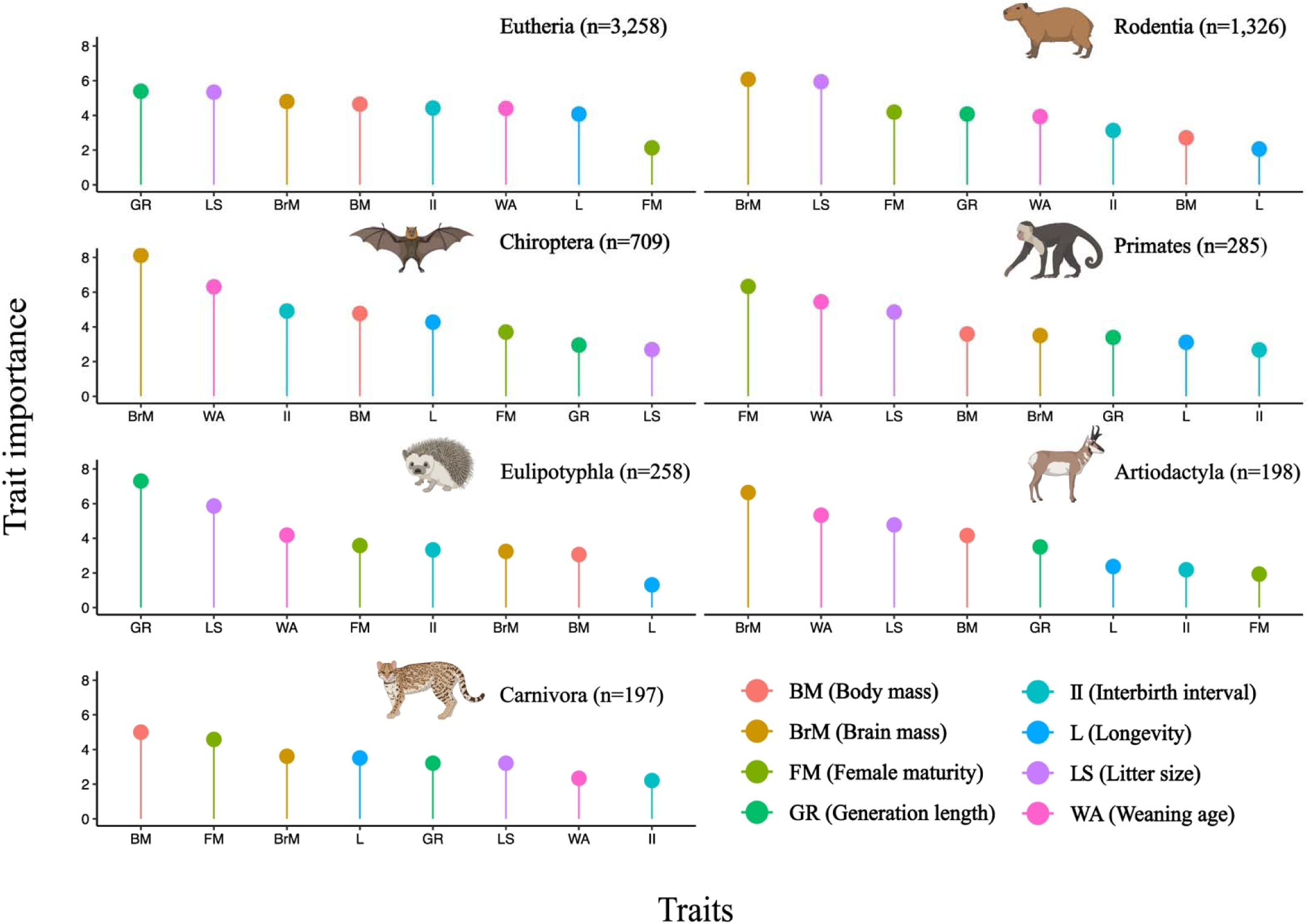
Variation in key predictive traits for gestation length across mammalian orders. The plots show the importance scores of eight traits in random forest analyses of predicting gestation length in all eutherian mammals (top left) as well as across six eutherian mammalian orders: Rodentia, Chiroptera, Primates, Eulipotyphla, Artiodactyla (excluding Cetacea), and Carnivora (excluding Pinnipedia). The x-axis of each plot depicts the eight traits in order of their importance scores (from highest to lowest) in predicting gestation length, and the y-axis shows the actual importance scores. Each colored marker (lollipop) in these plots represents a specific trait, with the height of the lollipop indicating its importance score. The analyzed traits included Body Mass (BM), Brain Mass (BrM), Female Maturity (FM), Generation Length (GR), Interbirth Interval (II), Litter Size (LS), Longevity (L), and Weaning Age (WA). Silhouette illustrations were created in BioRender.com.

To further explore the evolutionary associations between gestation length and other eutherian traits, we performed phylogenetic regression of gestation length with all other traits as covariates using Pagel’s model [30] (see **Methods**). We found that body mass, relative brain mass, and weaning age were generally strongly positively correlated with gestation length, while litter size was generally strongly negatively correlated in multiple datasets (**Table S2**). Associations of gestation length with other traits varied by order. For example, longevity was significantly correlated only in Eulipotyphla and Chiroptera (p-values < 0.01), female maturity was generally not correlated except in Eulipotyphla (p = 0.04), and generation length was significantly correlated in Eutheria, Rodentia, and Eulipotyphla (p-values < 0.01). These patterns underscore the robustness of the association between gestation length and some traits (i.e., body mass, brain mass, litter size, and weaning age) across eutherian orders and the varying significance of others (i.e., longevity and generation time), emphasizing both the commonalities and the differences in the life history strategy patterns of different mammalian orders.

### Variation in gestation length drives variation in relative brain mass and litter size in eutherian mammals

To investigate potential causal relationships between gestation length and other traits, we performed phylogenetic path analysis on a set of 20 plausible hypotheses of how the nine traits interact with each other, paying particular attention in how gestation length interacts with the other eight traits [18,20,31] (see **Methods**). We identified various causal relationships between gestation length and the other eight traits across different mammalian orders (**Fig. 2**). Most notably, we found that gestation length causally influenced both relative brain mass (positively) and litter size (negatively) in Eutheria and individual orders with two exceptions. The first exception was the Primates, where the relationship between gestation length and relative brain mass was equivocal; the two best supported hypotheses were statistically indistinguishable, with one favoring a weak causal influence of gestation length on relative brain mass and with the other favoring the inverse relationship (**Fig. 2**). The second exception were the Chiroptera, where the causal influence of gestation length on litter size in Chiroptera was observable, albeit not significant. Additionally, in Primates, the relationships between relative brain mass and longevity, as well as interbirth interval with both weaning age and longevity, were also equivocal (see **Methods**). Moreover, gestation length causally influenced weaning age, although both the sign (positive/negative), and to some extent the strength (correlation coefficient), of this association varied across orders.

**Fig 2.**
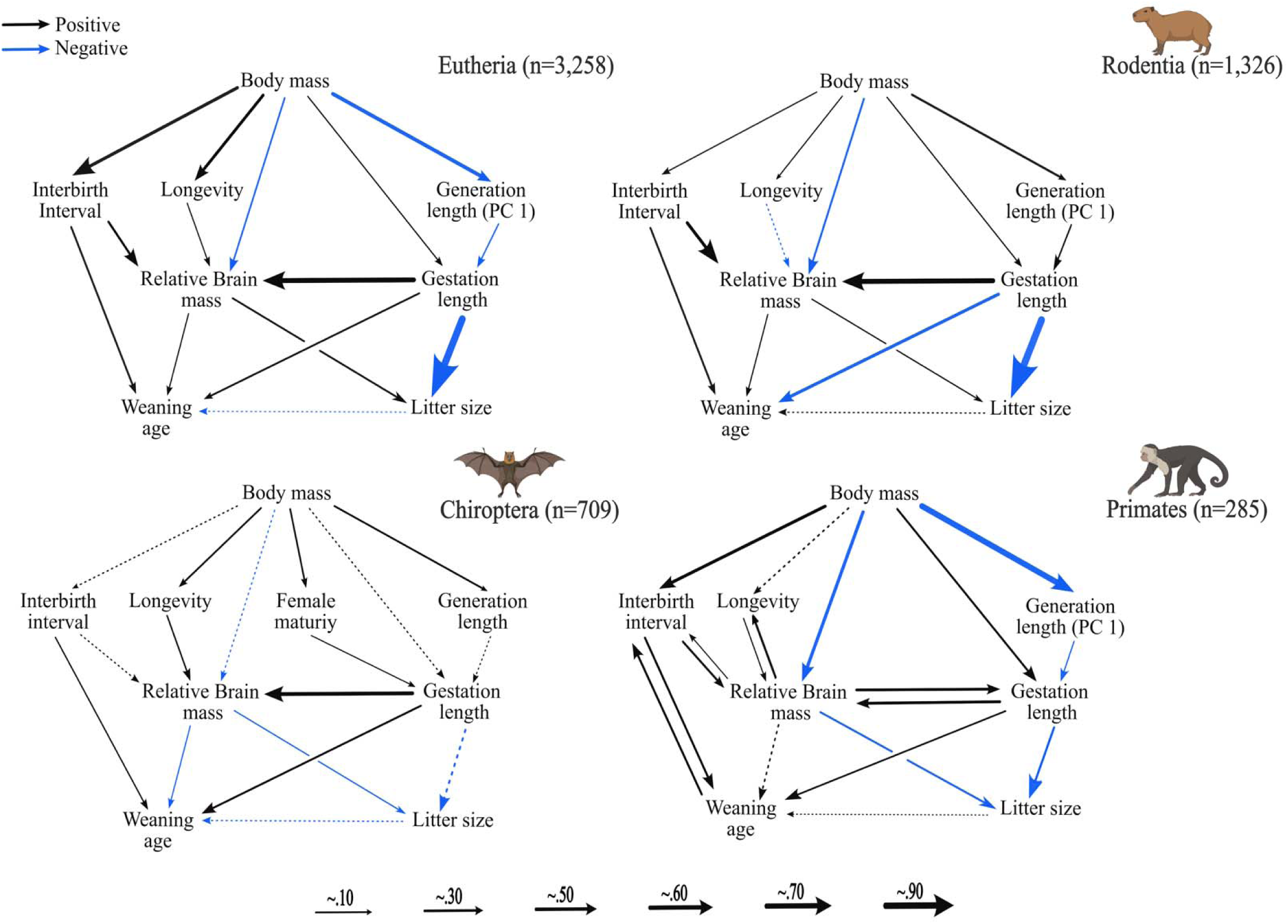
Gestation length variation drives variation in relative brain mass and litter size in eutherian mammals. This figure depicts the causal associations among various traits in Eutheria (n=3,258), Rodentia (n=1,326), Chiroptera (n=709), and Primates (n=285), assessed through d-separation phylogenetic path analyses of 20 different plausible hypotheses of trait relationships [18–20] (see **Fig. S1** for graphs for the rest of the orders; see **Figs. S2 and S3** for all the hypotheses tested in our path analysis). In Primates, bidirectional arrows represent that the top two hypotheses were statistically indistinguishable with one hypothesis favoring trait-trait association in a particular direction (e.g., gestation length causally influences relative brain mass) and the other hypothesis favoring the opposite direction (e.g., relative brain mass causally influences gestation length). PC 1 represents the main component capturing the largest variance between generation length and female maturity, reducing the multicollinearity of those two traits, in Eutheria, Rodentia, and Primates (see **Methods**). Arrow thickness represents the strength of the correlation coefficients, with thicker arrows indicating stronger associations. Black arrows indicate significant positive correlations, while blue arrows indicate negative correlations. Dashed arrows represent non-significant correlations. Silhouette illustrations were created in BioRender.com.

There is a substantial body of literature on the allometric relationship of gestation length with body mass in eutherian mammals [1,2,5,6,8,9,11–13,32,33], which has given rise to the conventional notion that body mass is a main driver in the evolution of gestation length. Interestingly, we found that the causal phylogenetic association between body mass and gestation length was not significant in Chiroptera and Eulipotyphla and weakly significant in Eutheria and Rodentia (r = 0.21). In contrast, we observed a moderate significant association in Primates (r = 0.27), Artiodactyla (r = 0.30), and Carnivora (r = 0.40).

These results suggest that gestation length directly influences brain mass and litter size across eutherian mammals and is directly influenced by body mass or other traits only in select orders.

### Gestation length is strongly coevolving with brain mass (positively) and litter size (negatively)

The phylogenetic path analysis [18–20] revealed that gestation length directly influences brain mass and litter size across eutherian mammal orders. We also observed causal relationships between gestation length and body mass and weaning age in several orders. To further explore evolutionary correlations between gestation length and these four traits, we quantified their strengths using the ratematrix [22] Bayesian framework (see **Methods**).

We found that the evolutionary correlation patterns between gestation length and the four traits (**Fig. 3**) were strongly concordant with the predicted causal relationships (**Fig. 2**). Specifically, the strongest positive association among all pairs of traits was between gestation length and brain mass and the strongest negative association was between gestation length and litter size (**Fig. 3**). We also observed a moderate correlation between gestation length and both body mass and weaning age, in agreement with the weaker (and/or more variable between orders) causal relationships between gestation length and these two traits. The evolutionary correlation of gestation length and body mass is the weakest among the trait pairs studied and varies between orders (**Figs. 3 and S4**). For example, consistent with the results of the phylogenetic path analysis [18–20] (**Fig. 2**), gestation length and body mass are weakly negatively correlated in Chiroptera but strongly positively correlated in Primates and Carnivora (**Figs. 3 and S4**).

**Fig 3.**
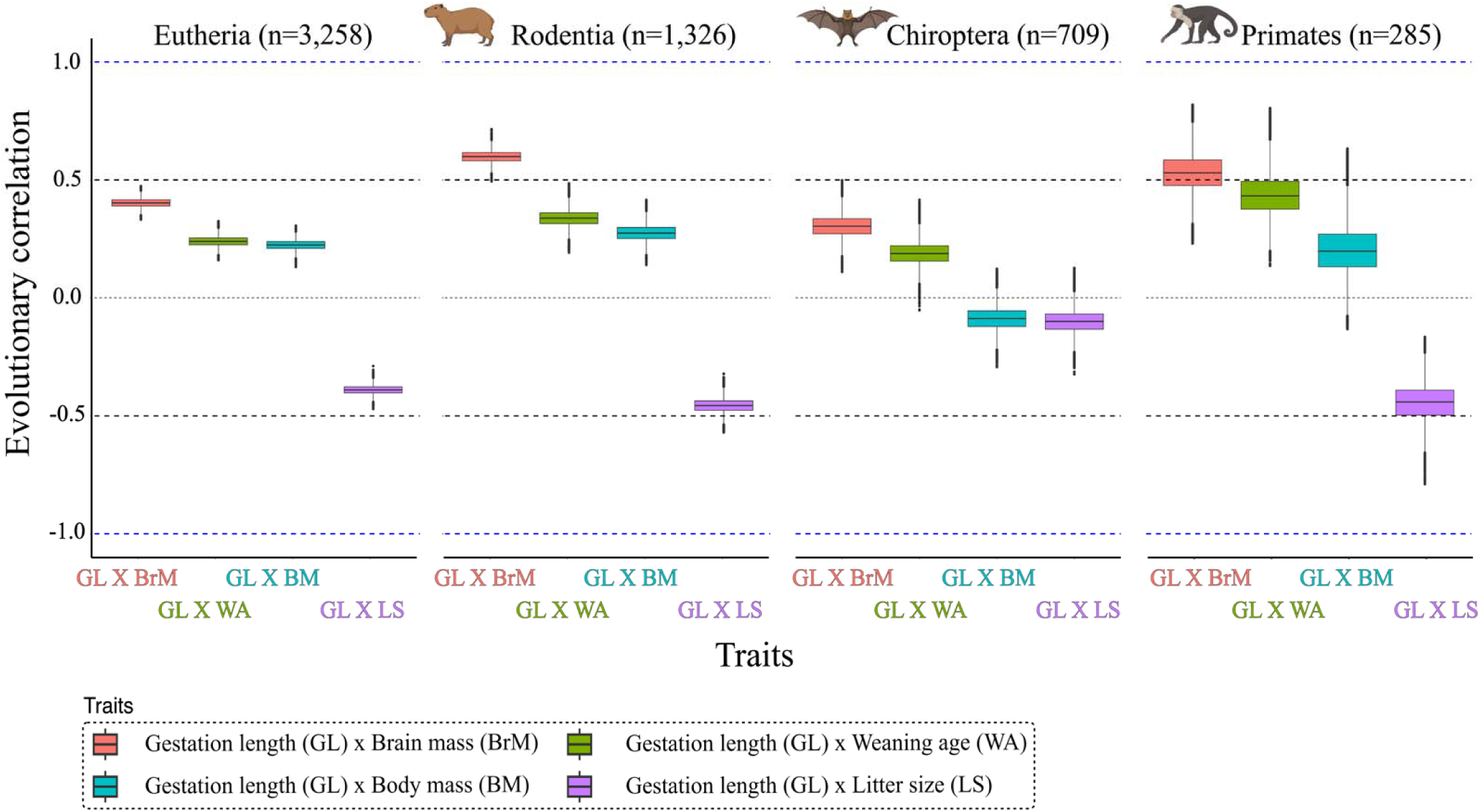
Strong evolutionary correlation between gestation length with brain mass and litter size in across mammals. This box plot shows the evolutionary correlation between gestation length and four key traits (brain mass, litter size, weaning age, and body mass) across four mammalian orders: Eutheria (n=3,258), Rodentia (1,326), Chiroptera (n=709), and Primates (n=285). The x-axis of the plot depicts the different trait pairs, and the y-axis shows the Bayesian estimation of evolutionary correlation scores for each pair of traits derived from ratematrix [22] analy is. Each color represents a different pair of correlated traits, highlighting the variation and strength of trait correlations across the groups displayed in the different panels. Silhouette illustrations were created in BioRender.com.

## Discussion

The major question addressed in this study concerns the causal relationships between gestation length and key life history traits, as well as other important traits, in eutherian mammals. Our findings suggest that longer gestation periods in eutherian mammals support increased brain development (**Fig. 2**). Considering the substantial energy demands associated with brain development [34–38], this prolonged gestation presumably facilitates further prenatal brain growth and development. Additionally, we found that gestation length negatively and causally influences litter size, suggesting that species with longer gestation periods tend to produce smaller litters. This inverse relationship likely results from the increased maternal investment required for extended prenatal development, limiting the number of offspring a mother can support during gestation [13,39,40]. Species that invest in fewer offspring (longer gestation periods) give birth to more developed young, who are better equipped to survive in resource-scarce or high-pressure environments [41–43]. In contrast, species that produce larger litters often have shorter gestations, relying on quantity to ascertain fitness [13,44]. This reflects a broader spectrum in evolutionary trade-offs where species with slower life histories—characterized by longer gestations and fewer, more developed offspring—balance reproductive output with offspring quality and survival [13,39].

The energetic demands of brain development have long been recognized as a key factor shaping gestation length across mammals and other animals [2,4,14,45–50]. Because developing and maintaining large brains is energetically costly [35,51–53], the evolution of gestation length is closely tied to these energetic demands. Species with longer generation lengths can allocate more resources to individual offspring, facilitating growth of larger neonate brains [54]. Our inference that gestation length variation causally and positively influences brain size variation suggests that gestation length plays a critical role in mammal life history patterns (**Fig. 2**) [2,4,45–47]. This causal link could also reflect trade-offs between life history strategies favoring slow development and extended parental care, requiring longer gestation for sufficient brain development before birth, versus strategies favoring fast development and limited parental care [2,16].

The path analysis [18–20] revealed that gestation length strongly and negatively affects litter size in most mammalian orders (**Fig. 2**). Species with longer gestations prioritize fewer, more developed offspring, in contrast to species with shorter gestation periods that produce larger litters. This suggests that species with longer gestation periods tend to have smaller litters, likely because extended prenatal development requires more substantial maternal investment in each offspring (altricial species), limiting the number of viable fetuses [45,55–58]. While longer gestation periods can benefit offspring survival and development in some cases, such as in the precocial guinea pigs and spiny mice [5,57–61], postnatal factors such as maternal behaviors (e.g., infanticide, siblicide) and environmental pressures can significantly influence the effective litter size [43,55,62]. Additionally, maternal physiology, ecological conditions, and species-specific adaptations can all play a role in determining litter size and reproductive strategies [55]. These factors often create energetic constraints during development, especially when cognitive abilities critical for energy acquisition are still maturing.

While the strong correlation between gestation length and body mass when the two traits are examined in isolation is well established [1,10,12,33,63], our analyses suggest that this relationship is weaker and more complex. There are two explanations for our findings. First, prior studies often rely on smaller sample sizes disproportionately influenced by well-studied orders, such as primates, artiodactyls, and carnivores, which represent only a fraction of eutherian diversity. Second, the inclusion of several additional traits in our analyses, which like gestation length are also typically correlated with body mass [1,4,6,12,33,39,64–66], revealed that some of them play a substantial role in shaping eutherian mammal gestation length. This suggests that when the correlation between gestation length and body mass is analyzed in isolation, some of the variation explained by these other traits is falsely attributed to body mass. Our findings of a causal link between body mass and gestation length only in certain orders and its absence in others emphasize that gestation length is a multifaceted trait influenced by multiple interacting factors beyond body mass.

The pattern of causal relationships between gestation length and other traits in Primates differs from the patterns observed in other orders (**Fig. 2**). For example, we identified equivocal links between brain mass and gestation length, a unique effect not observed in the other orders studied. A previous study of 46 primates found a direct association between gestation length and brain mass, suggesting that gestation length influences brain mass [67]. Conversely, Sacher and Staffeldt [2] reported that brain mass imposes a constraint on gestation length. Our analyses could not disentangle the direction of causality in the relationship between primate gestation length and brain mass, which are likely shaped by a combination of ecological pressures, social structures, and metabolic demands influencing primate evolution as well as by differences within primate clades [45,68]. This complexity underscores the need for further research to clarify how the life history strategies of different primate clades differ from each other and from those other mammals.

Our study clarifies the complex causal relationships between gestation length, brain mass, litter size, and body mass, emphasizing the central role of gestation length in shaping mammal life histories. This work highlights the intricate network of interdependencies between these traits, offering valuable new insights. Nevertheless, several caveats must be acknowledged. While we included a broad array of traits to capture a diverse set of species, important factors such as neonate developmental status, brain mass, and metabolic rate were not included due to limited data availability. Incorporating these traits could provide a more comprehensive understanding of the causal relationships, potentially revealing an even stronger influence of gestation length on neonate brain mass. Moreover, the lack of fossil data, particularly for life history traits, hinders our ability to precisely time when specific traits evolved, which limits the reconstruction of historical relationships between gestation length and other traits. Recent advances, such as extracting life history information from mammalian fossil teeth [69], offer exciting opportunities to address this gap and enhance our understanding of evolutionary drivers. Integrating such methods could further clarify the processes shaping the causal links between life history traits. Finally, path analysis [18–20], by design, involves selective modeling which restricts our ability to exhaustively examine every potential influence. As new empirical data become available, future studies should incorporate this knowledge to refine our understanding of evolutionary shifts in gestation length and related traits.

## Methods

### Data collection

We obtained data on gestation length (GL) and eight other mammalian traits: body mass (BM), brain mass (BrM), female maturity (FM), weaning age (WA), interbirth interval (II), generation length (GR), litter size (LS), and longevity (L) for 3,258 eutherian mammal species from publicly available databases, including PanTHERIA [24], AnAge [25], EltonTraits [26] and COMBINE [27]. To ensure data quality, we included only adult values for each species and conducted a thorough manual inspection. Prior to analysis, we natural log-transformed all continuous predictor variables to reduce skewness. We used the mammalian phylogeny inferred by Álvarez-Carretero et al. [23], ensuring that all analyses were conducted on mammal species that matched the phylogeny for consistency. This comprehensive framework enabled us to account for evolutionary relationships among the species in our analysis. Cetacea and Pinnipedia were excluded due to their unique aquatic adaptations, which significantly impact their life history traits and physiology, and small sample sizes [28,29]. This exclusion helps prevent skewed results and maintains focus on the evolution of gestation length in terrestrial mammals. The dataset used in this study is provided in **Data S1**.

### Trait selection with Random Forest

We employed Random Forest (RF), using the randomForest [21] function for feature selection to identify the most important predictors from the collected datasets. This effectively managed multicollinearity and provided insights into the relative importance of each feature in predicting gestation length. We log-transformed and scaled the variables across datasets. Following preprocessing, a random forest model was applied using k-fold cross-validation (k=10) to assess model performance across different hyperparameters, including the number of trees (50, 100, 200), tree depth (5, 10, 15), and thresholds for variable importance (0, 0.01, 0.03, 0.05). Feature importance was calculated based on the %IncMSE metric [70].

### Phylogenetic regression

All statistical analyses were carried out in the R 4.3.3 [71]. Due to the strong correlation between body mass and brain mass in mammals [4,72,73], we used relative brain mass, i.e., brain mass scaled by body mass, to increase the explanatory power of our analyses. In addition, in certain datasets (Eutheria, Rodentia, Primates), the age of female maturity and interbirth interval were strongly collinear in our models based on the Variance Inflation Factor (VIF) measure, which we calculated using the function ‘check_collinearity’ from the sensiPhy [74] R-package (**Table S2**). In cases of collinearity, we used Principal component analysis (PCA) [75] to reduce these to one uncorrelated component (Eutheria, Rodentia and Primates). Based on its loadings, the first principal component describes female maturity (**Table S2**), so in our analysis PC 1 reflects the trait female maturity. We verified that none of the other traits exhibited significant collinearity; all VIF values were < 2.5, after incorporating the first PC and accounting for phylogeny **(Table S2**).

Comparisons among species, due to their shared ancestry, violate the assumption that trait values are independently drawn by a common distribution [76]. To account for this lack of independence, we employed phylogenetic generalized least squares regression analyses (PGLS) using the ‘phylolm’ function in the R package phylolm [77] and Pagel’s λ [30] model. We analyzed the datasets using the following models:

For Chiroptera, Artiodactyla, Eulipotyphla, and Carnivora:

log(gestation length) ∼ log(longevity) + log(body mass) + residuals(log(brain mass) ∼ log(body mass)) + log(interbirth interval) + log(generation length) + log(litter size) + log(weaning age) + log(female maturity).

For Eutheria, Rodentia and Primates:

log(gestation length) ∼ log(longevity) + log(body mass) + residuals(log(brain mass) ∼ log(body mass)) + log(interbirth interval) + log(litter size) + log(weaning age) + log(female maturity x generation length)(PC 1).

### Phylogenetic path analysis

We conducted phylogenetic path analyses [18–20] to explore the causal evolutionary relationships between gestation length and the other eight traits. We tested 20 distinct models, each of which represents a plausible hypothesis of how gestation length causally influences or is influenced by the other eight traits. Each model was defined by a set of d-separation statements [20], representing hypotheses through a directed acyclic graph linking all predictors with assumed causal relationships. The selection of these graphs and models was informed by our random forest and phylogenetic regression analyses and was based on two criteria (**Fig. S2-S3**). First, we kept the established link between body mass and brain mass (body mass influencing brain mass) constant across all models. Second, we selected models that enabled us to test hypotheses about causal relationships between gestation length and the eight other traits. Specifically, we tested models where body mass, female maturity, and generation length potentially influence gestation length, while gestation length potentially influences brain mass and litter size. Additionally, we explored models where brain size might be a causal factor for gestation length. Each of the 20 models tested was ranked on its C-statistic information criterion (CICc) using the R package phylopath [20]. In cases where the ΔCICc > 2, we used the top model [18–20], otherwise we averaged the top models, using the “average” function from phylopath package [20].

### Evolutionary correlation analysis

To test the evolutionary correlation between mammalian traits, we used the R package ratematrix [22,78]. All trait data were transformed using the natural logarithm. We ran two independent Markov Chain Monte Carlo (MCMC) chains in parallel, each consisting of 500,000 generations, with a burn-in proportion of 40%. We set a log-normal prior on the evolutionary variances of all traits, with both the mean and standard deviation set to 0.1. We checked the acceptance ratios for both chains and tested for convergence between them. Upon achieving convergence, we merged the two MCMC chains and plotted the rate matrix to assess the evolutionary correlation between the traits.

## Data availability statement

All data, scripts, and supplementary information associated with this manuscript will be made publicly available upon acceptance for publication.

## Conflict of interest statement

A.R. is a scientific consultant for LifeMine Therapeutics, Inc. The authors have no other conflicts of interest.

