## Supplementary Figures for "Gestation length drives brain size and litter size variation in eutherian mammals"

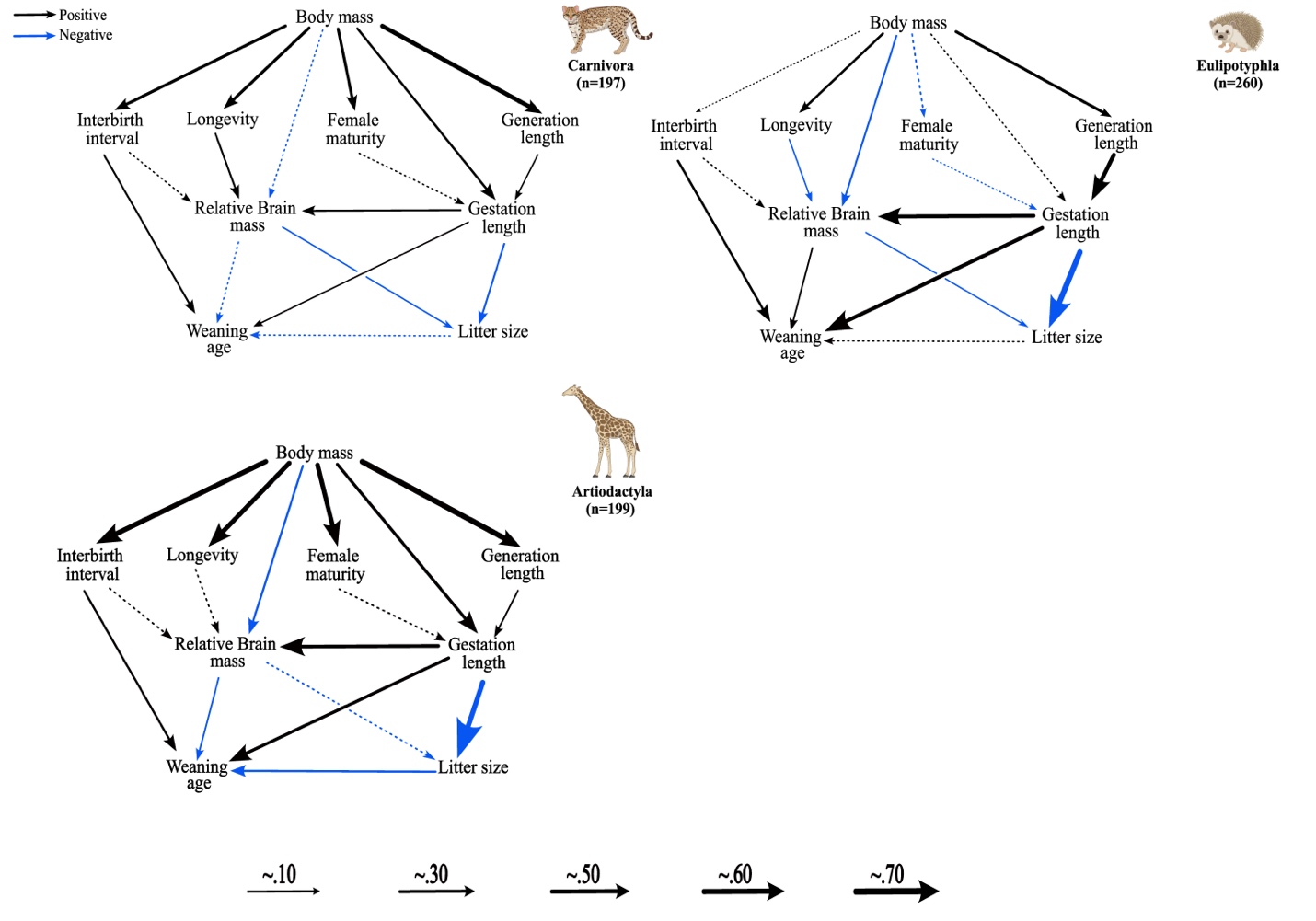

**Fig S1. Gestation length variation drives variation in relative brain mass and litter size in eutherian mammals.**

This figure depicts the causal associations among various traits in Carnivora (n=197), Eulipotyphla (n=260), Artiodactyla (n=199), assessed through d-separation phylogenetic path analyses [1]. Arrow thickness represents the strength of the correlation coefficients, with thicker arrows indicating stronger associations. Black arrows indicate significant positive correlations, while blue arrows indicate negative correlations. Dashed arrows represent non-significant correlations. Silhouette illustrations were created on Biorender.com.

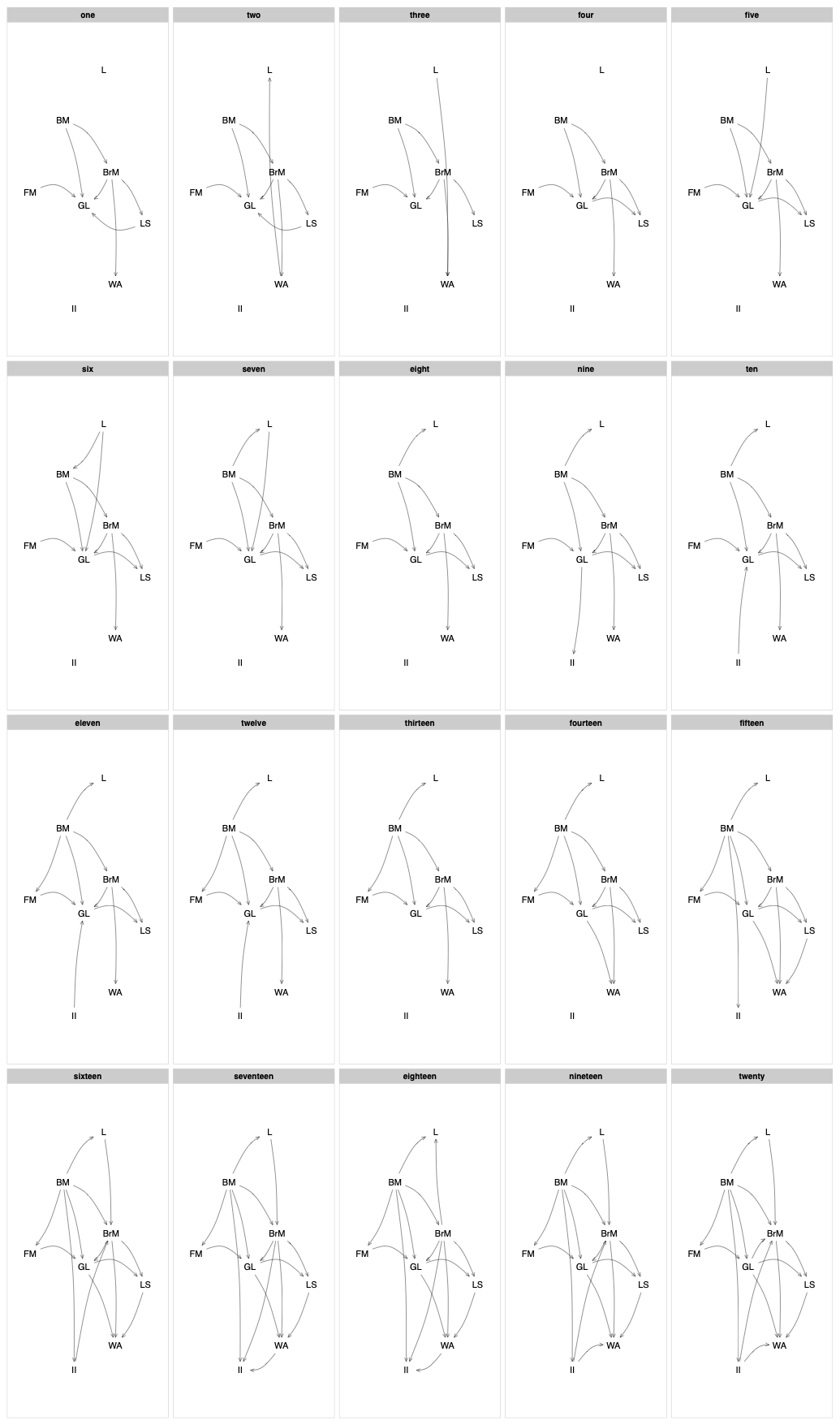

**Fig S2. Directed acyclic graphs depicting the path models evaluated in our phylogenetic path analysis.**

Directed acyclic graphs showing the 20 models tested in the phylogenetic path analysis of Carnivora (n=197), Eulipotyphla (n=260), Artiodactyla (n=199), and Chiroptera (n=709). The analyzed traits included Body Mass (BM), Brain Mass (BrM), Female Maturity (FM), Generation Length (GR), Interbirth Interval (II), Litter Size (LS), Gestation Length, Longevity (L), and Weaning Age (WA).

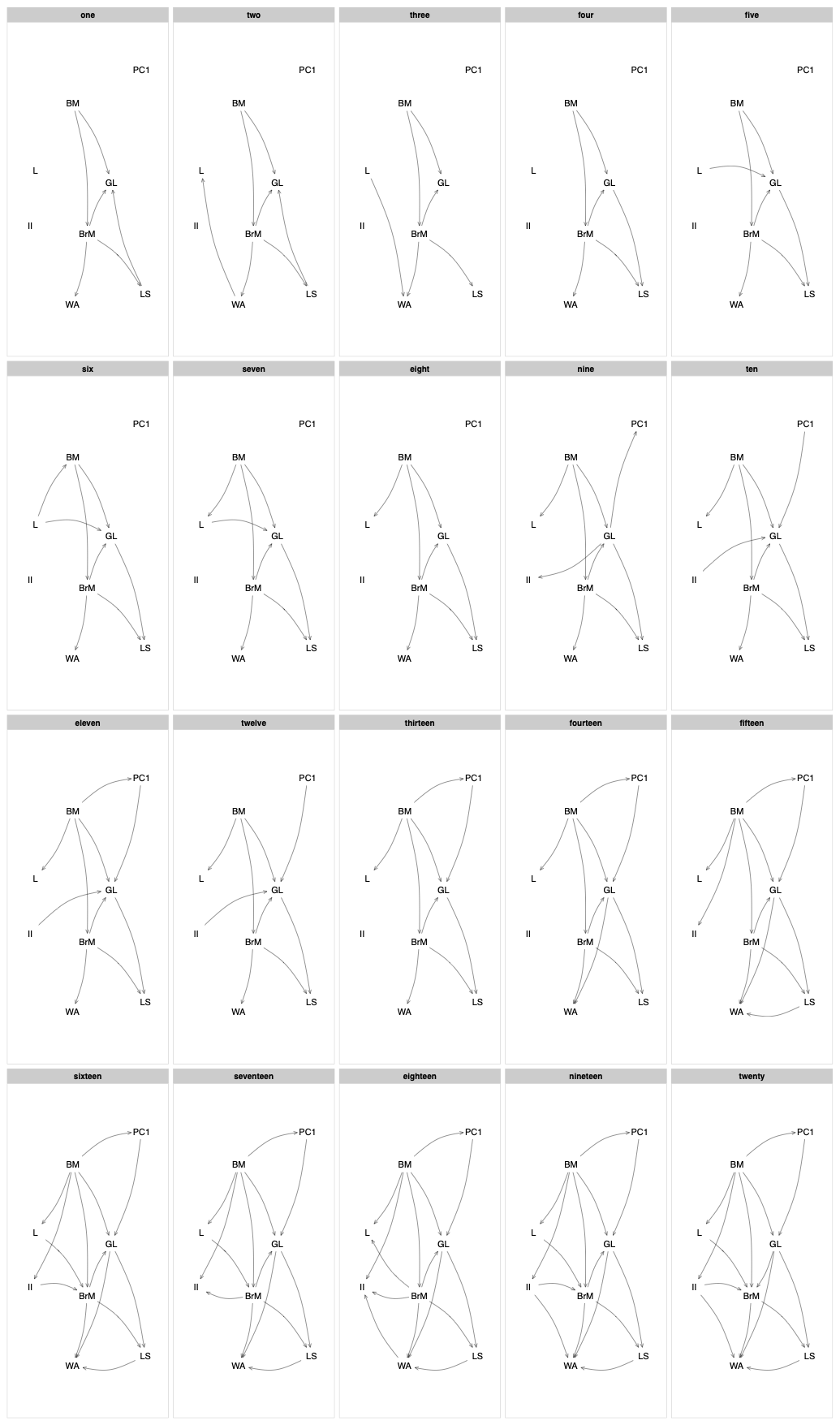

**Fig S3. Directed acyclic graphs depicting the path models evaluated in our phylogenetic path analysis.**

Directed acyclic graphs showing the 20 models tested in the phylogenetic path analysis of Eutheria (n=3,258), Rodentia (n=1,326), and Primates (n=285). The analyzed traits included Body Mass (BM), Brain Mass (BrM), PC 1 (Generation Length x Female Maturity), Interbirth Interval (II), Litter Size (LS), Gestation Length, Longevity (L), and Weaning Age (WA).

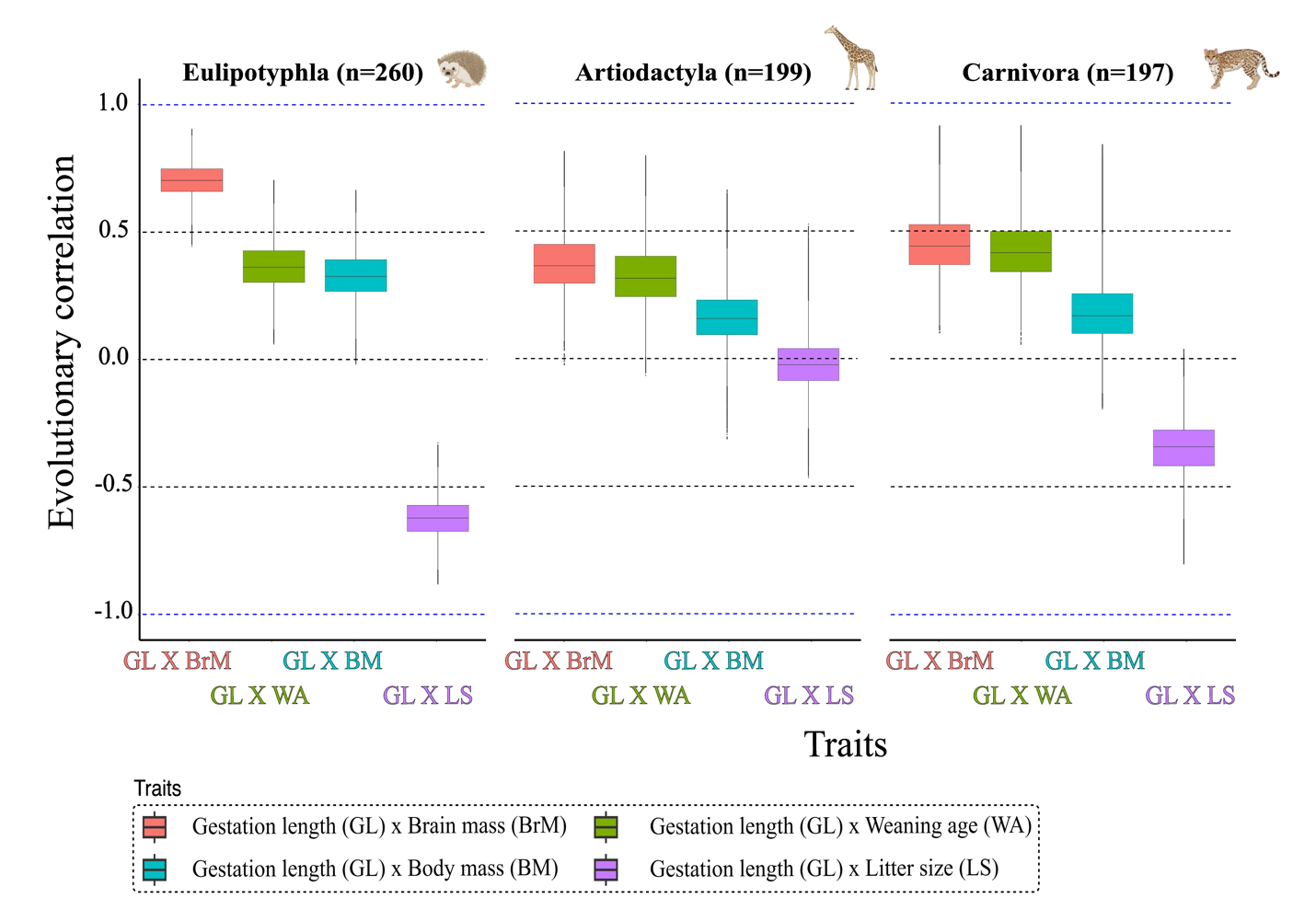

**Fig S4. Strong evolutionary correlation between gestation length with brain mass and litter size in across mammals.**

This box plot shows the evolutionary correlation between key traits across three mammalian orders: Eulipotyphla (n=260), Artiodactyla (199), and Carnivora (n=197). The x-axis of the plot depicts the different traits, and the y-axis shows the Bayesian estimation of evolutionary correlation scores for each pair of traits derived from ratematrix [3] analysis. Each color represents a different order, highlighting the variation and strength of trait correlations within each group. Silhouette illustrations were created on Biorender.com.
